# Interplay between folding and binding modulates protein sequences, structures, functions and regulation

**DOI:** 10.1101/211524

**Authors:** Bálint Mészáros, László Dobson, Erzsébet Fichó, Gábor E. Tusnády, Zsuzsanna Dosztányi, István Simon

**Affiliations:** MTA-ELTE Momentum Bioinformatics Research Group, Department of Biochemistry, Eötvös Loránd University, Pázmány Péter stny 1/c, Budapest, H-1117 Hungary; Protein Structure Research Group, Institute of Enzymology, RCNS, Hungarian Academy of Sciences, Magyar Tudósok krt 2, Budapest, H-1117 Hungary; Membrane Protein Bioinformatics Research Group, Institute of Enzymology, RCNS, HAS, Budapest PO Box 7, H-1518, Hungary

## Abstract

Intrinsically Disordered Proteins (IDPs) fulfill critical biological roles without having the potential to fold on their own. While lacking inherent structure, the majority of IDPs do reach a folded state via interaction with a protein partner, presenting a deep entanglement of the folding and binding process. Protein disorder has been recognized as a major determinant in several properties of proteins; yet the way the binding process is reflected in these features in general lacks this detail of description. Recent advances in database development enabled us to identify three basic scenarios of the interplay between folding and binding in unprecedented detail. These scenarios have fundamentally different properties in terms of protein sequence, structure, function and regulation, depending on the structural properties of the interacting partners. Strikingly, the existence of a binding partner and its structural properties influence all analyzed properties of proteins to the same extent as the divide between inherent order or disorder. The appreciation of this interplay between folding and binding is the basis for the successful charting of unknown territories in the protein interactome, the understanding of how different binding modes assemble regulatory networks, and the development of future pharmaceutical applications.

## Introduction

Proteins deliver the basic machinery for life, providing functions indispensable for all living organisms. The foundation of the molecular understanding of how proteins function was hallmarked by the determination of the first protein structures. The resulting dogma, called structure-function paradigm (Redfern et al., 2008) delineated the central thesis of structural biology: protein function is born from structure, and the prerequisite of a functional protein is prior successful and complete folding.

In the late 90's mounting evidence led to the realization that ordered proteins, which conform to the dogma, represent only part of the protein world (Wright and Dyson, 1999). There are several other functional proteins that lack a stable tertiary structure in their isolated form. Although they defy previous dogma, intrinsically disordered proteins (IDPs) are critically important functionally, especially in signaling and regulation (Dyson and Wright, 2005; Wright and Dyson, 2015). With this birth of ‘unstructural biology’ (Tompa, 2011), the protein world was divided into two major regions. This binary view is deeply embedded in us at the conceptual level, exemplified by current disorder prediction methods (Deng et al., 2015; Dosztányi et al., 2010) and their evaluation schemes (Monastyrskyy et al., 2014). As an extension of this binary representation, it has been long recognized that protein flexibility is rather a continuous spectrum, ranging from (almost) true random coils (Gast et al., 1995), through molten globules (Sutovsky and Gazit, 2004) and proteins that are marginally stable (Wang et al., 2013), to stable domains.

A description of IDPs complementary to their flexibility stems from the consideration of their interactions. While some IDPs stay disordered while exerting their function - some even when bound to a protein partner (Tompa and Fuxreiter, 2008) -, the vast majority of known IDPs do adopt a stable conformation as a result of interacting with a partner protein. In these cases the folding happens at the same time as the binding, and the two processes, governed by the same biophysical forces (Tsai et al., 1999), are deeply intertwined.

The entanglement of folding and binding for IDPs results in binding modes clearly distinct from interactions of ordered proteins. The interaction between an IDP and an ordered protein partner – termed *coupled folding and binding* (Dyson and Wright, 2002) – holds the potential for forming weaker, transient interactions due to the loss of conformational entropy decreasing the binding strength (Chu and Wang, 2014). The study of the specific structural properties of such interactions gave rise to a better understanding of this binding mode (Mészáros et al., 2007), enabled targeted prediction development (Malhis et al., 2016; Meng et al., 2017; Mészáros et al., 2009), and ultimately led to successful development of novel ways of pharmaceutical modulation through the development of small molecules (Shen and Maki, 2011).

In contrast, complexes formed exclusively by IDPs – through a process termed *mutual synergistic folding* – are far less understood. Most of our knowledge stems from individually studied cases (Demarest et al., 2002) and analyses of relatively small datasets (Gunasekaran et al., 2004; Rumfeldt et al., 2008). While several related classes of protein complexes have been analyzed (such as intertwined complexes (Mackinnon et al., 2013)), these works define their focus interactions based on the properties of the bound structure instead of the structural states of the unbound proteins. However, this lack of targeted analysis of mutual synergistic folding is primarily due to the lack of data, as until recently no databases existed focusing on various types of IDP interactions in structural detail (Fichó et al., 2017; Schad et al., 2017).

In this work we assess the basic types of interaction between proteins, considering both ordered and IDP interactors. We pioneer how the sequence properties of the binding sites, the structure of the resulting complexes, the fulfilled biological roles, and the regulation of the interactions reflect the way participating proteins reach an ordered structure: on their own, or via interaction with an ordered or an IDP partner. Using this information we are able to get a full view of the entire spectrum of protein-protein interactions for the first time. Furthermore, we are able to explore the intrinsic classes of complexes formed via mutual synergistic folding rooted in sequence and structural properties. This presents the first classification system paralleling ones existing for ordered proteins (Andreeva et al., 2008; Pearl et al., 2003).

## Results

### 1. Interplay between folding and binding is reflected in the amino acid composition

Four protein interaction categories were considered in the analysis based on how constituent protein chains reach a structured state. These include proteins going through autonomous folding and independent binding, coupled folding and binding, mutual synergistic folding, and IDPs presumably not forming interactions and hence not adopting a structure.

Figure 1 and Table S1 show the calculated sequence properties for each group. In terms of sequence composition, interacting ordered proteins on average resemble closely the reference residue composition of the human proteome, with a marked decrease in prolines, incompatible with ordered secondary structures. Intrinsically disordered regions (IDRs) not involved in protein-protein interactions (PPIs) conform to the generic view of the typical residue composition of disordered proteins - i.e. depleted in stabilizing residues and enriched in structure breaking residues (Campen et al., 2008). In contrast, the sequence compositions of interacting IDRs show distinct differences, strongly reflecting the structural state of their binding partner. IDRs recognizing ordered proteins are often highly charged, lack hydrophobic residues, and often contain prolines. On the other hand, IDRs binding to other IDRs are typically more hydrophobic (on par with ordered proteins) and contain very few prolines/glycines (even less than the average for ordered proteins). They are also often highly charged and devoid of cysteines and aromatic residues, highlighting the markedly diminished role of disulfide bridges and π stacking in their structure formation. Although there are clear trends discriminating the four groups, variances within residue groups are high, reflecting the heterogeneous nature of proteins in all structural types.

**Figure 1:**
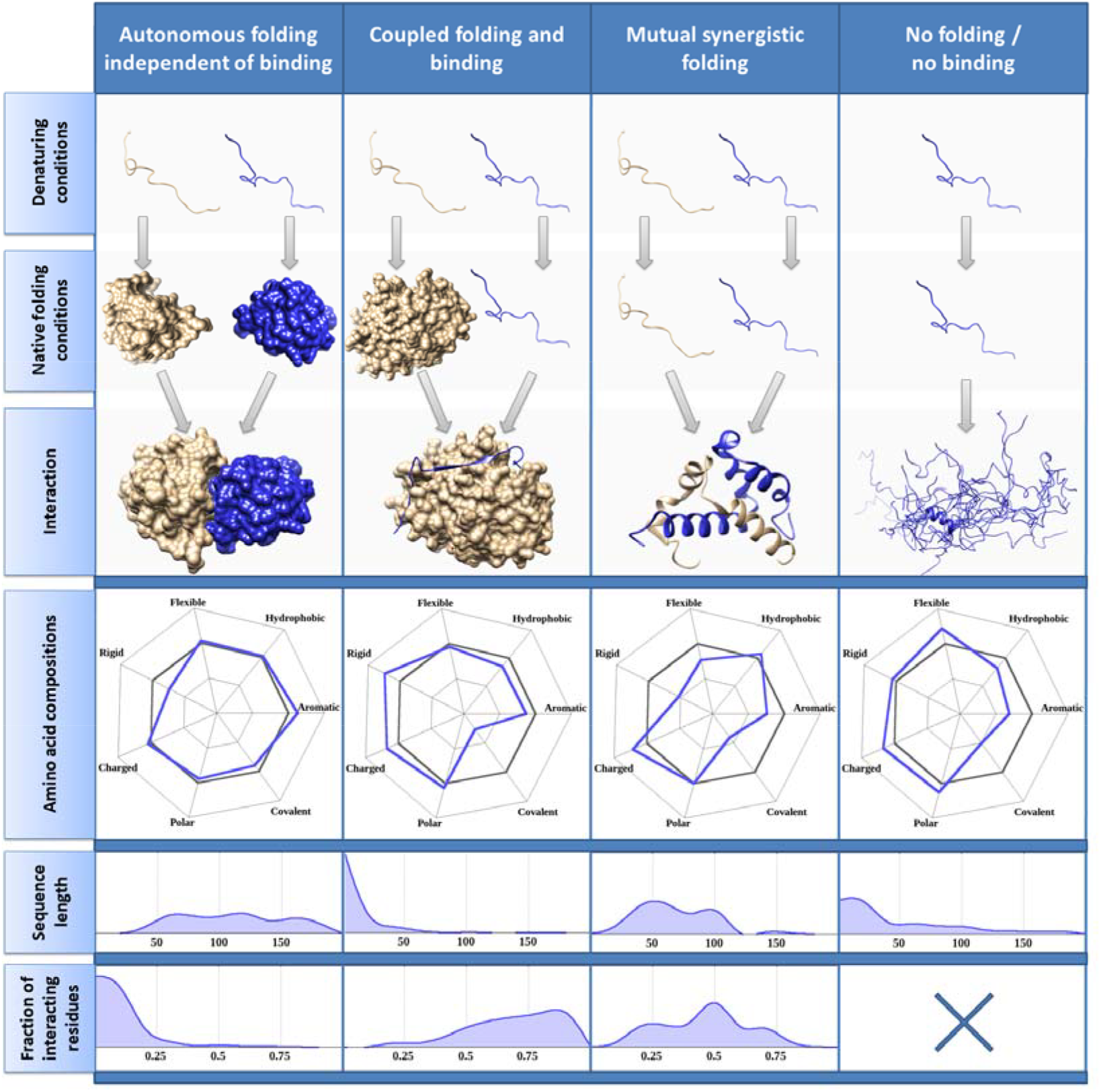
Sequence properties of proteins based on the relationship between their folding and binding. Columns mark the four basic ways a protein can reach a structured state. Radar charts show the relative amino acid content compared to the human proteome (grey). Amino acids are grouped according to their biochemical/structural properties: hydrophobic (A, I, L, M, V), aromatic (F, W, Y), polar (N, Q, S, T), charged (H, K, R, D, E), rigid (P), flexible (G), and covalently interacting (C). Amino acid groups on the right and left side of the radars represent residues commonly considered to be stabilizing and structure breaking, respectively (Campen et al., 2008). The lowermost two panels contain the sequence length distribution and the fraction of residues directly involved in the interaction, respectively.

Regarding sequence lengths, interacting ordered proteins on average contain more residues as the presence of a folded domain in incompatible with extremely short sequences. Non-interacting IDPs and IDRs mediating interactions with ordered proteins tend to be significantly shorter, while proteins with mutual synergistic folding are on par with ordered proteins in terms of sequence length. Taking into consideration the fraction of residues directly involved in the interactions with the partner reveals a new layer of distinctive features. Ordered proteins use only a low fraction of their residues in the interaction due to sterical reasons. In contrast, IDPs tend to donate a larger relative number of residues to the binding, with several IDRs undergoing coupled folding and binding consisting entirely of interacting residues.

The uncovered characteristic differences in terms of sequence properties imply different binding modes for the three studied interacting groups. The differences between the length and the interacting content of the affected protein regions hint at basic structural differences, motivating a deeper structural analysis of the bound structures.

### 2. The presence of protein disorder modulates structural properties of the bound conformation

The structures IDPs and ordered proteins adopt upon binding to a partner were analyzed (see Data and Methods), with a focus on secondary structures, surface areas, atomic contacts and predicted interaction energies (Figure 2, Table S2).

**Figure 2:**
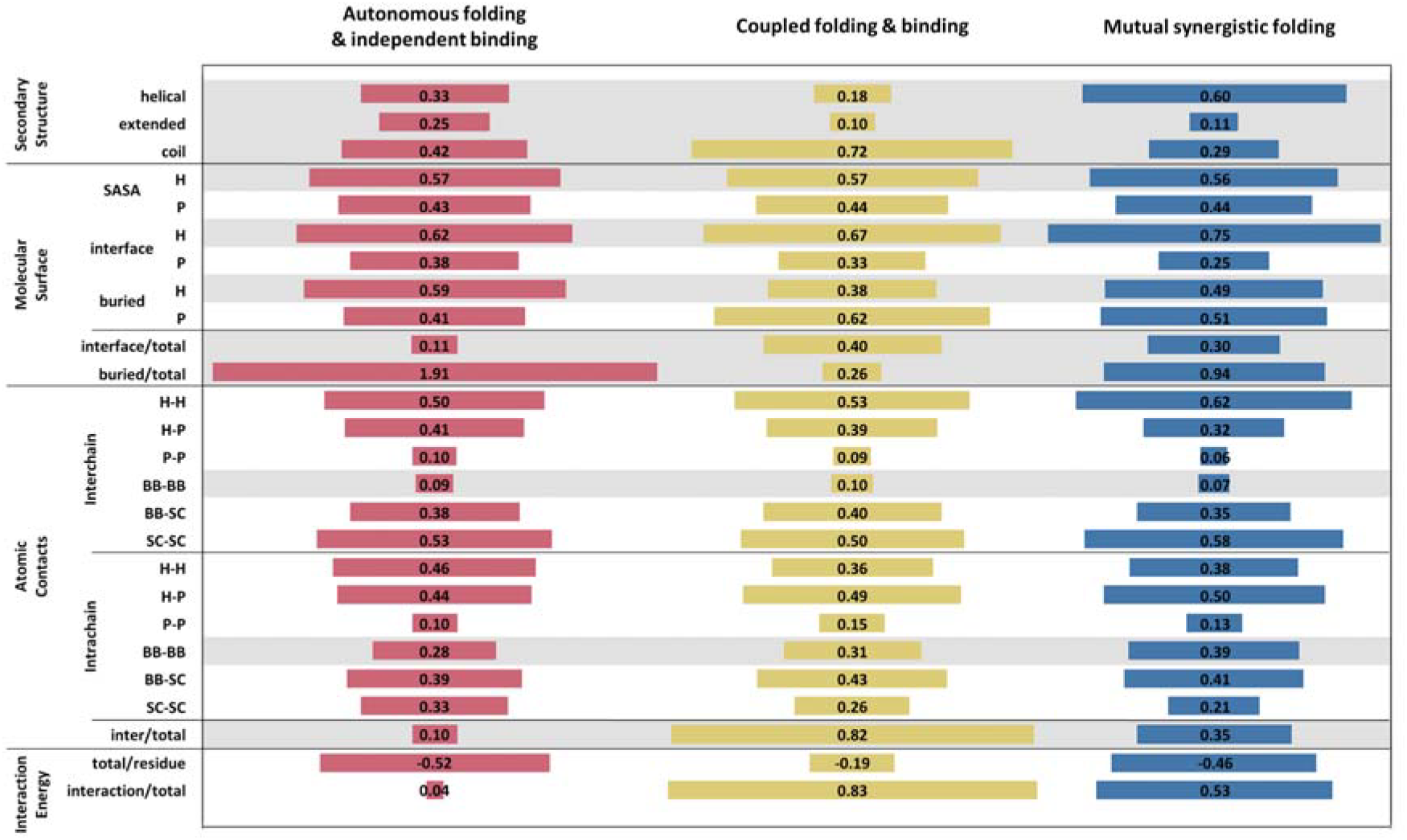
Normalized average structural properties of proteins as a function of their folding and binding process. Columns mark the different interaction groups. Grey background highlights weakly correlated features. Abbreviations: SASA: Solvent Accessible Surface Area; H: Hydrophobic; P: Polar; Bb: Backbone; Sc: Sidechain.

Ordered proteins show a relatively balanced composition of both helical and extended secondary structures, and residues outside periodical secondary structures connecting them. Compared to this balanced structural makeup, bound IDP structures show pronounced differences. IDPs folding on the surface of ordered proteins generally lack ordered secondary structures and adopt irregular structures. On the contrary, IDPs undergoing mutual synergistic folding show a strikingly strong preference for helical structures.

Molecular surface areas primarily describe the hydrophobic effect with the surrounding solvent (Richmond, 1984). In the bound form, all three types of proteins have similar hydrophobic/polar (H/P) ratio of solvent accessible surface areas - they all exist in the same aqueous environment. However, their interfaces are highly different with hydrophobicity playing a more important role for the binding of IDPs. In contrast to this shielding effect of the partner, polar surfaces are typically buried in IDPs, i.e. they are made inaccessible due to intramolecular shielding.

The relative sizes of different molecular surfaces are also highly distinctive. IDPs on average utilize a much higher fraction of their molecular surfaces in interactions compared to that of ordered proteins. Complexes formed exclusively by IDPs retain a considerable fraction of their surfaces as solvent accessible, while IDPs binding to and folding on the surface of ordered proteins use by far the largest fraction of their available surfaces as interaction interfaces. Buried surfaces show an inverted trend with IDPs burying only a small fraction of their surface when bound to ordered partners, in contrast to synergistically folding IDPs and ordered proteins.

In terms of atomic contacts, interactions between hydrophobic atoms aids interchain interactions, while hydrophobic-polar contacts play a major role in intrachain interactions and this trend is more pronounced for IDPs. Interchain interactions are primarily mediated through side chains. Intrachain interactions, however, are evenly formed by sidechain and backbone atoms in the case of ordered proteins. In contrast, for IDPs backbone atoms play a clearly more important role. The ratio of interchain and intrachain contacts clearly shows that IDPs utilize their residues more efficiently for binding the partner protein. As the sequence of IDPs undergoing coupled folding and binding is usually shorter (see Figure 1), a biologically meaningful stability has to be established by a small number of interacting residues. Although IDPs with mutual synergistic folding display a larger number of intrachain interactions, they are still more heavily dominated by the interaction with the partner, compared to ordered proteins.

The properties and relative extents of calculated surfaces and contacts all contribute to the overall stability of the resulting complex. In order to assess this stability, interaction energies were calculated based on residue-level statistical potentials (see Data and Methods). According to the energy calculations, ordered protein complexes are the most tightly bound systems on average. Synergistic folding of IDPs result in comparable stabilizing per residue energies, however, the per residue stabilizing energy of IDPs bound to ordered proteins is significantly weaker, possibly corresponding to the prevalence of more transient interactions. The relative energetic weight of the interaction between subunits in the overall stability is low for ordered complexes, but over ten- and twenty-fold higher for mutually folding IDPs and IDPs interacting with ordered proteins, respectively.

### 3. Various interactions mediate different biological functions with differential localization

It is known that the functional repertoire of IDPs in general is distinctively different from that of ordered proteins (Wright and Dyson, 2015). We extended the study of protein functions by analyzing the characteristic processes conveyed by the three types of interactions. Functional annotations were based on Gene Ontology (GO) terms describing biological processes (see Data and Methods).

In order to make GO annotations at highly different levels directly comparable, a reduced version of the ontology (PPI GO Slim – see Data and Methods and Table S3) was created containing a limited set of higher-level terms covering a wide range of possible biological functions; and all original GO annotations were mapped to the PPI GO Slim. The most commonly occurring PPI GO Slim terms for all three classes of interactions are shown in Figure 3. Generic high-level processes, such as communication, transport and development, are executed via a large number of carefully coordinated interactions from all three interaction classes. However, processes involving a more restricted number of interactions show specificity towards interaction types, such as immune responses and the maintenance of homeostasis for ordered complexes, or cell division, cell cycle control, cell adhesion, and viral processes for interactions formed by IDPs.

**Figure 3:**
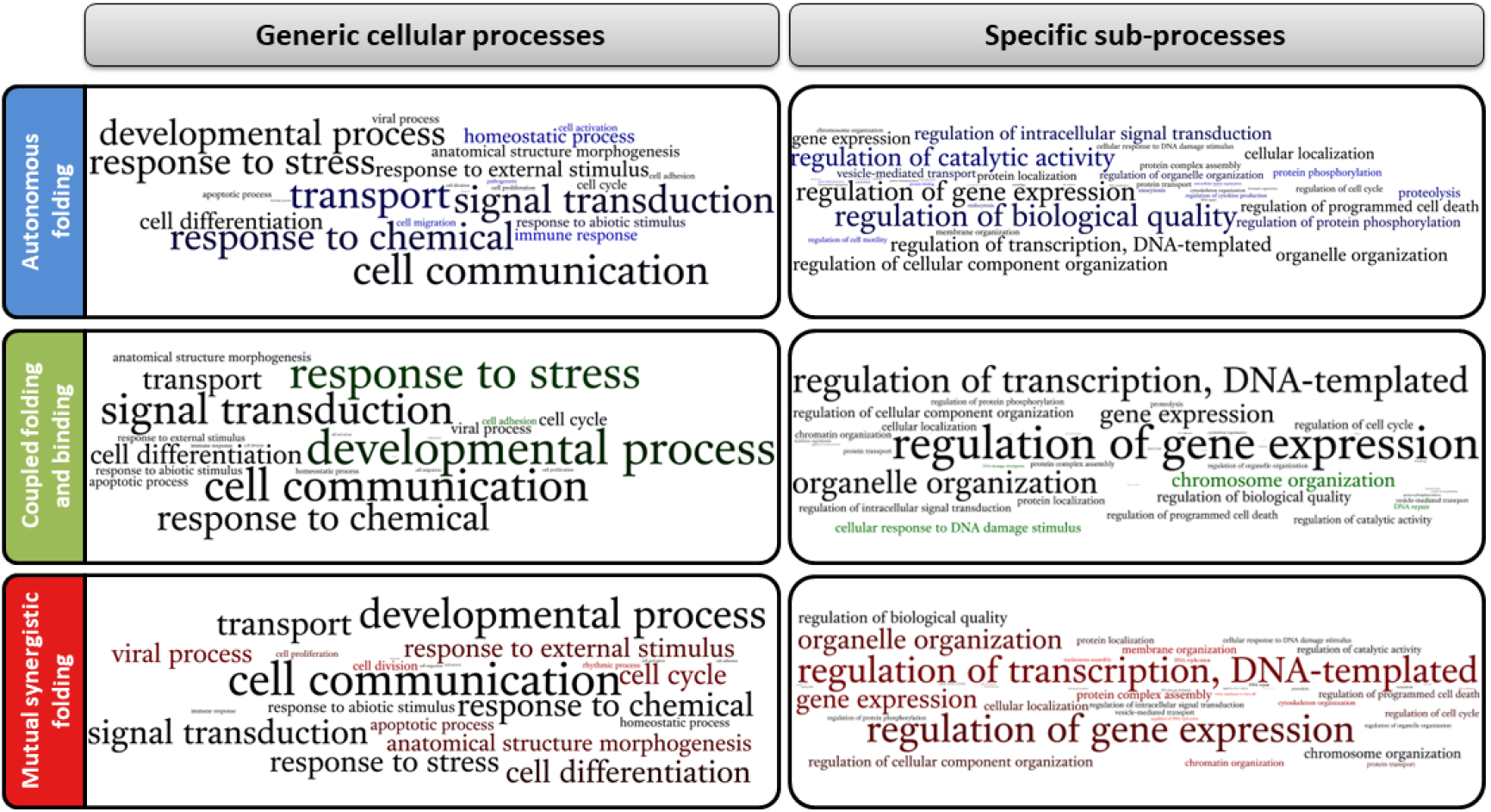
The frequency of occurrence for PPI GO Slim terms for the three classes of interactions. Left: generic terms, right: specific terms. Font size for a given biological process represents the relative frequency of occurrence of given GO term. Color depth represents the specificity of the term for the given interaction class. Process names in black show terms that ubiquitously occur in all classes, terms in full color represent terms that are unique to a type of interaction.

The analysis of specific sub-processes shows more pronounced distinctions between different interactions. The most prominent functions conveyed by ordered proteins are various regulatory processes, including the control of catalysis, biological quality, signal transduction and gene expression. Main IDP-mediated functions are primarily centered around DNA-processes, however there is a separation of functions depending on the structural state of the partner. Functions connected to the information storage function of DNA (often involving direct DNA contact), such as transcription and gene expression are clearly more characteristic of interactions formed exclusively by IDPs through mutual synergistic folding. On the other hand, processes pertaining to the regulation of DNA as a macromolecule, such as DNA damage response or chromosome organization, are dominated by IDP-ordered protein interactions.

GO annotations were also used to assess the typical subcellular localization of various interaction types (Figure 4) via CellLoc GO Slim (see Data and Methods and Table S3). The cytosol harbors a wide range of interactions from all types. Ordered interactions dominate the extracellular space and receptors embedded in membranes. Localizations closer to the DNA are progressively more dominated by IDPs: the nucleoplasm is the characteristic location for IDP-ordered protein interactions, and localizations directly connected to the DNA, such as the DNA packaging complex or the chromatin are the prime domains of mutual synergistic folding. Other common characteristic places of IDP-mediated interactions are non-membrane-bounded organelles, such as stress granules or the centrosome, falling in line with the recently realized importance of IDPs in the organization of such cell constituents (Boeynaems et al., 2017; Brangwynne et al., 2015).

**Figure 4:**
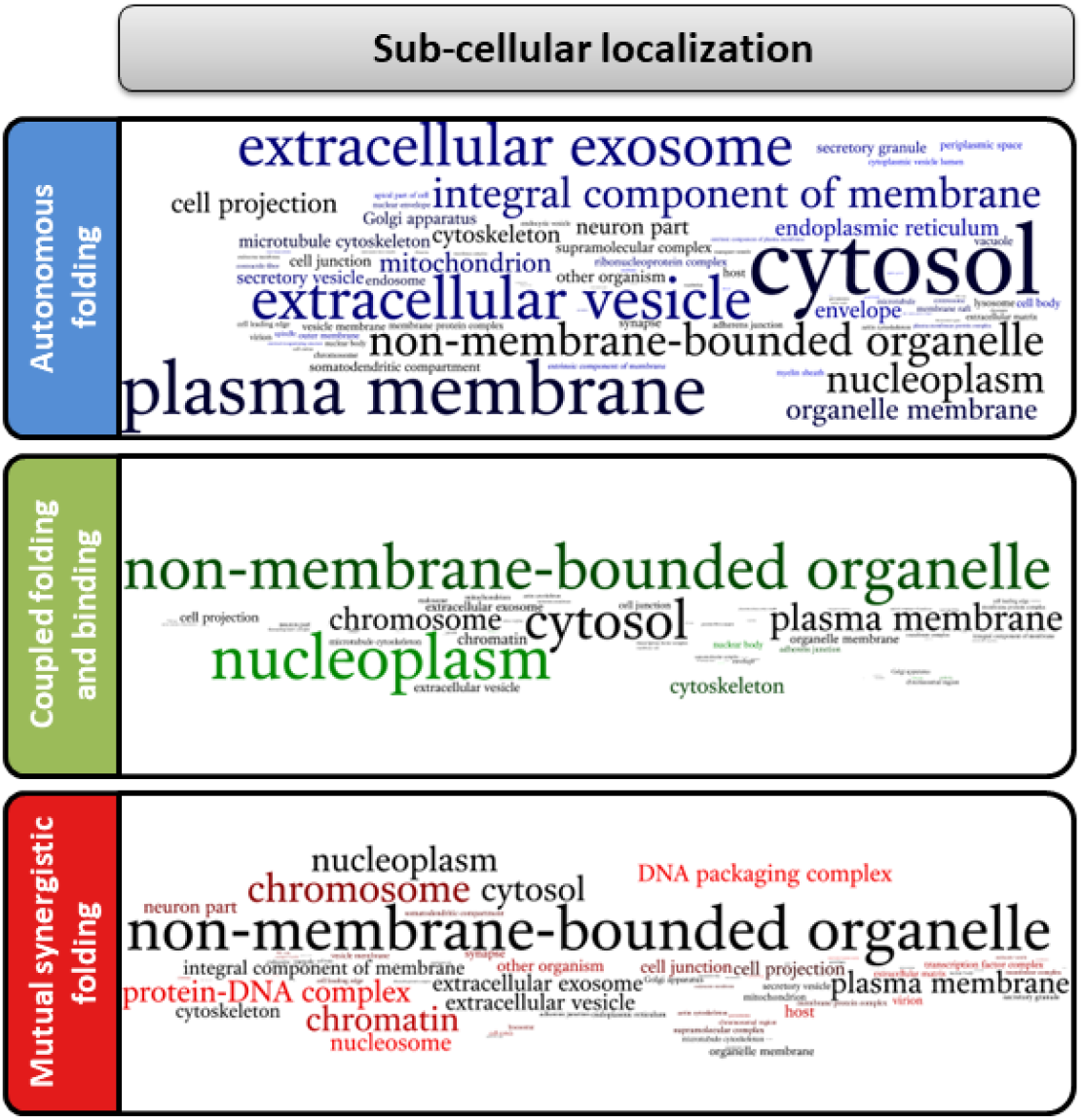
Characteristic sub-cellular localizations of the three classes of interactions. Font sizes represent the frequency of occurrence and font color depth marks the specificity of a cellular compartment for the given interaction type, similarly to the representation in Fig. 3.

### 4. Protein disorder modulates the heterogeneity of bound complex structures

The previous analyses have shown that interacting IDPs have distinctively differing residue compositions compared to interacting ordered proteins, and this composition reflects the structural state of the binding partner (Figure 1). In addition, the presence of IDPs in protein interactions heavily modulates the bound structure of resulting complexes and the functions these interactions mediate. However, these analyses only considered the average values of features, without quantifying the sequential, structural and functional heterogeneity of each interaction class.

In order to assess this heterogeneity, we first aimed to directly quantify and visualize the regions various protein complexes cover from the available sequence, structure and function spaces. These three levels were evaluated separately for all three interaction classes, taking into account the annotations of proteins described in the previous chapters (see Data and Methods). To visualize how proteins and interactions belonging to the three studied interaction classes are distributed in the sequence/structure/function space, Principal Component Analysis (PCA) was employed (see Data and Methods). Figure 5A shows the best two dimensional representation of these spaces, using the first two components carrying the highest fraction of variations of the data. While these variations are low, the visual inspection of the first two components can highlight basic differences of interaction classes. Furthermore, the fact that a large number of features are needed to represent the full variation of the data (see Figure S2) justifies previous sequence and structure feature selections, as well as the construction of the PPI GO Slim.

**Figure 5:**
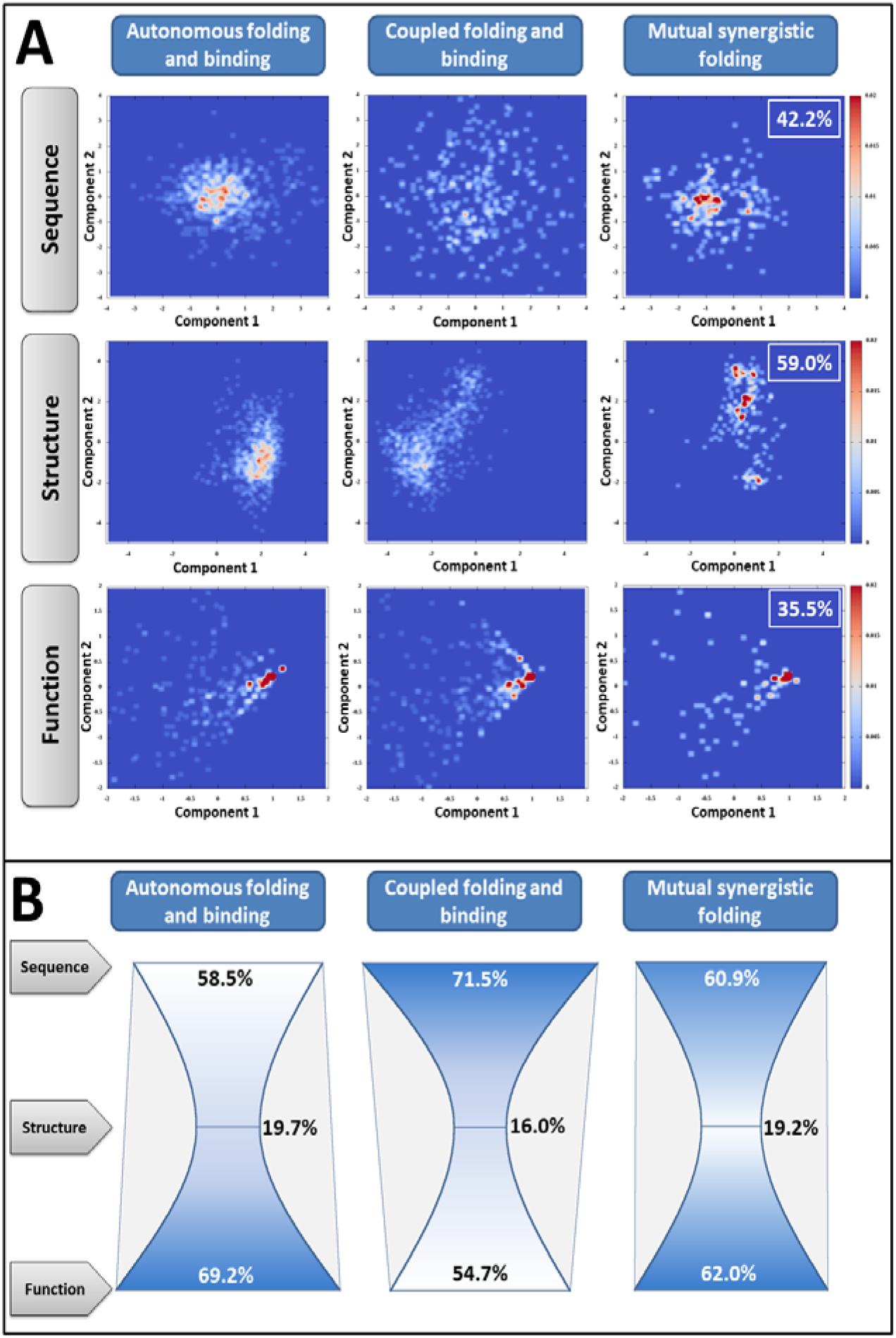
Variability of sequences, structures and functions for complexes from the three interaction classes. A: Distribution of various complexes considering the first two principal components of the sequence-, structure-, and functional space. Insets show the total variance of the data carried by the plotted components. B: Sequence-, structure-, and functional heterogeneity values calculated for all three classes of interactions.

Considering sequence-space distributions, complexes formed exclusively by ordered proteins or by IDPs shows a moderate demarcation, hinting at (at least partially) mutually exclusive residue compositions. IDPs capable of binding to ordered proteins, however, show a much wider distribution of compatible compositions, overlapping with both. In contrast, considering structural properties, all three classes seem to occupy a distinct subregion in the space of possible structures. Distribution of the three interaction types in the functional space shows a high degree of overlap. This reinforces the notion of the previous section stating that basically all high-level biological processes rely on both ordered proteins and IDPs as well, utilizing an interconnected network of their interactions.

To more objectively quantify the extent of various spaces used by different interactions, sequence-, structure-, and functional heterogeneity values were calculated for all three types of interactions. Heterogeneity values were defined as the average dissimilarity between two randomly chosen complexes from the same class. In turn, dissimilarity between two complexes from the same class was defined based on the hierarchical clustering of complexes (see Data and Methods). The so calculated heterogeneity values lie between 0% and 100%, with 0% corresponding to all complexes being identical and 100% corresponding to all complexes being as different as possible.

Calculated heterogeneity values (shown in Figure 5B) outline a basic trend. Regardless of structural state, proteins in general utilize highly variable sequence compositions to realize a comparatively much narrower set of structures. This reduction in complexity at the structure level, however, does not limit functional roles, as the functional heterogeneity of interacting proteins is on par with their sequential heterogeneity. Apart from general trends, the three classes of complexes show characteristic differences as well. Ordered proteins fulfill a wide range of functions with proteins of more restricted sequence compositions. IDPs undergoing coupled folding and binding represent the opposite using wide variations in composition, but conveying a more restricted range of functions in comparison. In contrast to both classes, complexes formed exclusively by IDPs show a striking balance between the heterogeneity of sequences used and the heterogeneity of biological functions they mediate.

### 5. Complexes of IDPs are tightly regulated at several levels

As shown in our previous functional analyses, all three studied classes of interactions play roles in crucial biological processes. In order for these processes to function correctly, the interactions on which they are built must be precisely regulated. These regulatory mechanisms include control of expression levels, subcellular localization, post-translational modifications, and competing interactions, among others. Previous studies have shown that IDPs are under exceptionally tight regulation (Gsponer et al., 2008), yet the interconnection between the structural state of the partner and the mechanisms of regulation is largely unknown. We analyzed three types of regulatory mechanisms for interactions: post-translational modifications (PTMs), alternative splicing, and competition between interactions.

Occurrences of four types of PTMs (phosphorylation, methylation, acetylation, and ubiquitination – see Data and Methods and Table S4) were studied for the three classes of interactions (see Figure 6A). All four types of PTMs are present on both ordered and disordered interacting proteins, with a pronounced accumulation of known PTM sites in IDPs undergoing mutual synergistic folding. In addition, these IDPs not only harbor more PTMs, but the occurrences of these modifications are far more correlated than for other proteins. This indicates a coordinated regulatory mechanism that is the most pronounced for complexes formed exclusively by IDPs.

**Figure 6:**
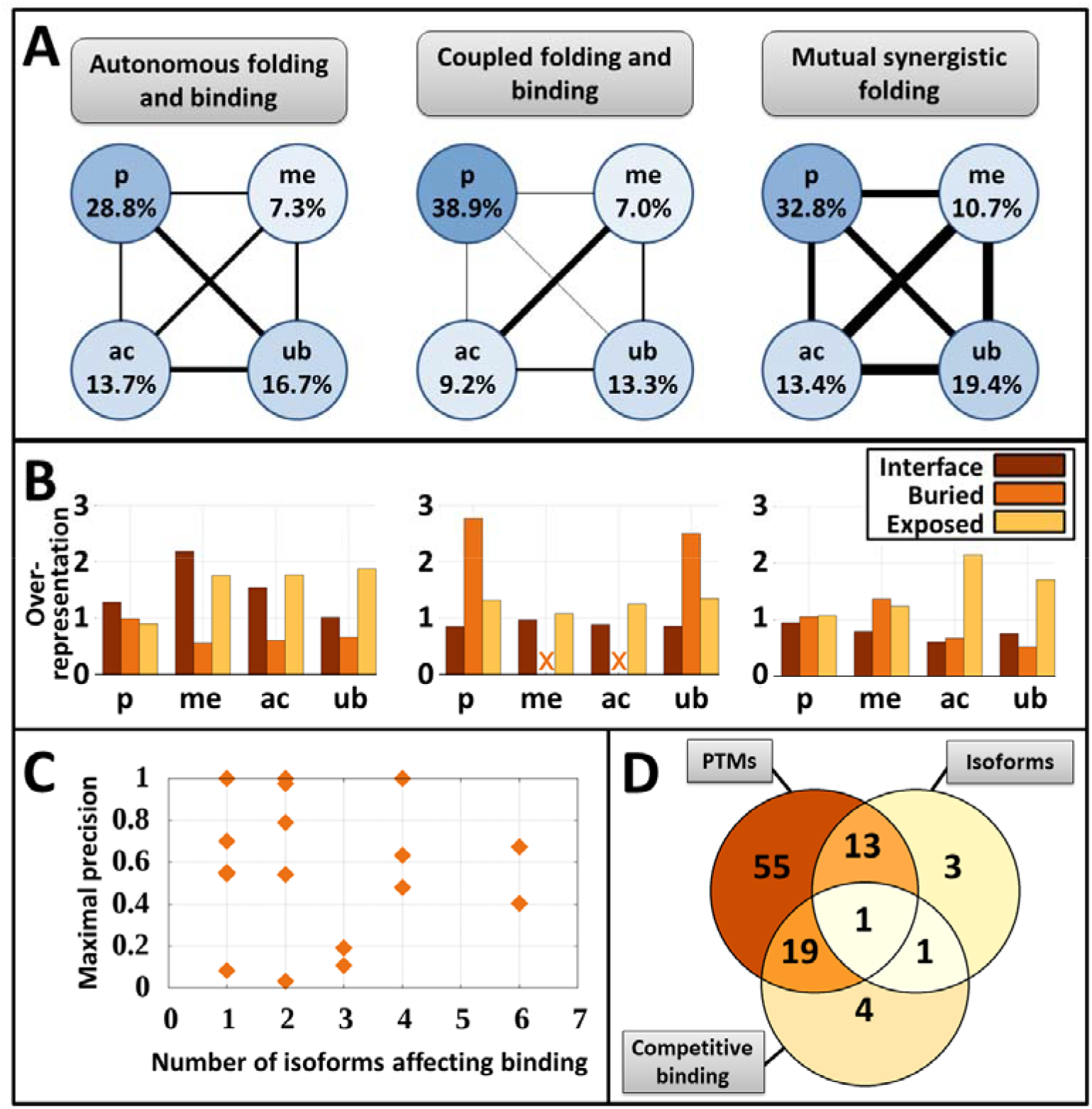
Regulatory mechanisms of interactions. A: the occurrence of PTMs in interacting proteins (p – phosphorylation, me – methylation, ac – acetylation, ub – ubiquitination). Color depth and percentage values represents the fraction of proteins affected. The width of connecting lines show the amount of mutual information between the occurrence of PTM pairs (see Table S4 for exact values). B: Location of PTM sites in the complex structure. Values represent enrichment compared to expected values based on the number of residues in each structural category. C: The number and specificity of isoforms affecting the binding regions of IDPs with mutual synergistic folding. Specificity represents the ratio of the spliced residues belonging to the binding site. For each protein only the specificity of the most specific isoform is shown. D: Number of IDPs undergoing mutual synergistic folding affected by the three types of regulatory mechanisms.

The structural location of PTMs offer insights into the mechanistic effects they have on the binding event (Figure 6B). Most ordered protein PTMs are enriched on the solvent accessible surface of domains, outside of the interface. These PTMs are not expected to directly modulate the binding, although they might have an indirect effect on the interaction (e.g. through controlling the availability of the protein via localization or degradation signals). As an exception, methylation and acetylation sites on ordered proteins show a tendency to preferentially target interface residues, directly modulating the interaction. In contrast, PTMs in IDPs generally seem to influence the binding event in a more indirect fashion. If the partner is an ordered protein, PTMs preferentially target buried residues (when there are any), sterically disrupting the native conformation the IDP would adopt upon binding. However, IDPs that bind to disordered partners are typically targeted through solvent accessible residues. As opposed to accessible residues of globular domains, these residues can heavily affect the binding through the tuning of local flexibility and predisposition for adopting a stable structure (Bah and Forman-Kay, 2016).

Alternative splicing is known to heavily affect short disordered binding regions binding to ordered partners (Buljan et al., 2013). Yet, the extent of control of mutual synergistic folding via protein isoforms lacks exploration. Figure 6C shows the number and precision of alternative splicing products for such IDPs where the splicing event directly affects the binding region. Most such proteins have only a few (typically 1-3) isoforms modulating the binding regions. Strikingly, in most of these cases, for at least one of the isoforms, the spliced protein region specifically targets the binding site. This indicates that alternative splicing not only affects synergistically folding IDP sites but specifically targets them.

The third common regulatory mechanism of interactions analyzed for complexes of IDPs is competitive binding. Mutually exclusive interactions are often observed for IDPs undergoing coupled folding and binding (Hsu et al., 2013; Weatheritt et al., 2012a). Our results (see Figure 6D and Data and Methods) show that many synergistically folding IDPs also have other known binding partners competing for the same binding region. Furthermore, the three types of interaction regulation are deeply intertwined with PTMs affecting both competitive binding and alternatively spliced regions. Interestingly, alternative splicing and competing interactions seem to present two alternative, largely disjoint mechanisms for interaction control. The only currently known example exhibiting all three types of regulatory mechanisms is the transactivation domain of p53.

### 6. Sequence and structure properties uncover the natural grouping of IDP complexes

Previous chapters have shown that all three interaction classes have distinctive sequential and structural features that on one hand enable their recognition as separate classes, and on the other hand show large enough variations to warrant the partitioning of these classes into specific subgroups. In case of ordered proteins, this grouping is rooted in their tertiary structures and has been addressed with various approaches of fold classification, such as SCOP (Andreeva et al., 2008) and CATH (Pearl et al., 2003). As IDPs undergoing coupled folding and binding bind to folded proteins, fold classes can serve as a basis of their structural classification as well (Schad et al., 2017). However, the classification of proteins going through mutual synergistic folding has not been tackled yet. The distribution of these proteins in the sequence-, structure-, and functional spaces (Figure 5) revealed, that structural features show the best potential of defining distinct groups, with additional information from sequential features. Functional features are distributed fairly evenly without showing trivial signs of a natural partitioning, and therefore are weak candidates to define inherent groups.

Sequential and structural features described in sections 1 and 2 were re-calculated for the full complexes of mutual synergistic folding (see Data and Methods). These features were used as input for hierarchical and k-means clustering to define 4 sequence-based and 5 structure-based clusters (Table S5). These sequence and cluster groups were manually compared and merged to provide a natural partitioning of synergistically folding IDPs into 6 groups, as shown in Figure 7. Apart from the main sequential and structural features, figure 7 also shows energetic properties, associated functions, subcellular localization, heterogeneity, regulatory mechanisms, and connection to grouping defined in MFIB.

**Figure 7:**
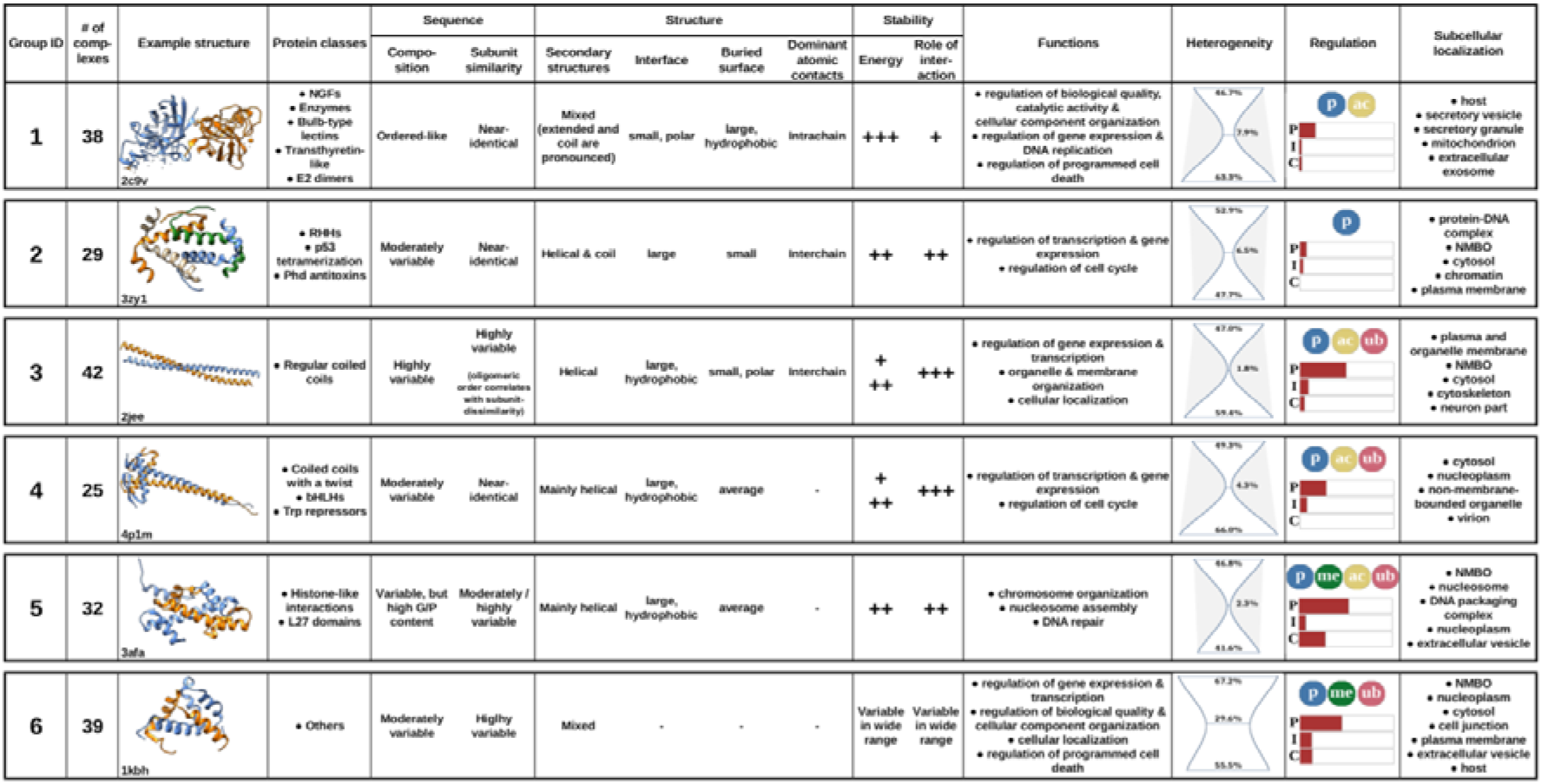
Groups of complexes formed by IDPs, based on sequence and structure features. NMBO-non-membrane bounded organelles. Sub-figures in the heterogeneity column show the group’s sequential, structural and functional heterogeneities, as calculated in section 4. Horizontal bars in the regulation column show the fraction of complexes in a given group involved in various types of regulatory mechanisms (P - post-translational modifications, I - isoforms affecting the binding regions, C - competing interactions). Color circles mark the dominant post-translational modifications for the group (p - phosphorylation, me - methylation, ac - acetylation, ub - ubiquitination).

The first group consists of complexes bearing a high similarity to ordered protein complexes. The constituent chains are highly similar, forming a large number of intrachain contacts, with inter-subunit interactions through a small polar interface playing only a secondary role in the stability of the complex. This group contains proteins with widely heterogeneous functions, including enzymes, transport proteins, and nerve growth factors. These proteins resemble ordered proteins in their localization as well, with extracellular regions being highly representative.

The second group also contains complexes with highly similar subunits. However, their sequence composition is distinctively different from that of ordered proteins, and instead of burying a large molecular surface to form a semi-stable hydrophobic core, they largely rely of the hydrophobic nature of their large interfaces. In accord, these structures are less bound and this stability depends more heavily of inter-subunit interactions. Interactions in this group play major roles in regulatory roles concerning transcription, gene expression and cell cycle, by including ribbon-helix-helix and prevent host death (Phd) antitoxin-type transcription factors, and members of the p53 family.

The third group is a structurally extremely homogenous protein complex family, composed entirely of coiled-coils. While their sequences are highly variable, their structures are exclusively composed of helices bound moderately/weakly through large hydrophobic interfaces. The role of interaction is even more pronounced in achieving stability compared to previous groups, as constituent proteins are able to bury only a small fraction of their polar surfaces. Coiled-coil interactions are often regulated, typically via various types of PTMs. Despite their highly similar structures, complexes in this group convey a large variety of functions, mainly pertaining to regulating transcription and performing membrane-associated biological roles, such as organelle and membrane organization.

Members of the fourth group are closely related to coiled-coils, however beside their classical helical interacting regions, they contain additional structural elements, e.g. a helix-loop-helix-type DNA interacting module. While the basic properties of group members closely resemble those of the previous group, subunits tend to be highly similar. Furthermore, interactions convey functions similar to those of group 2, focused on transcription, gene expression and cell cycle regulation.

The fifth group comprises complexes formed by different subunits moderately bound through a large hydrophobic interface. Complexes include the handshake fold of histone-like dimers and L27 domains. Group members are both structurally and functionally less diverse, with primarily DNA/chromosome-related functions. Interactions in this group are heavily regulated by all three studied mechanisms and all four kinds of PTMs.

While groups 1-5 represent well defined groups with members of evident similarities, the final group serves as an umbrella term for complexes that are not members of any previous structural/sequential classes. In accord, these complexes cannot be described by simple characteristic features and are the most sequentially and structurally heterogeneous group. This group contains highly specialized interactions that present unique protein complexes, which are highly regulated through all three control mechanisms.

## Discussion

Most current description of protein disorder treats this intrinsic structural property as either a binary feature or places proteins on a continuous spectrum of protein flexibility. However, even this extended classification considers only the unbound state of proteins. Our results clearly demonstrate that the capacity of binding to a partner protein and the structural state of that interactor has similarly deep consequences on the properties of IDP.

Possibly the most well-known distinguishing features of IDPs is their low hydrophobic content and high net charge, and this single observation opened the way for the construction of early disorder prediction methods (Uversky et al., 2000). In the past 15 years a more refined view of the sequential background of protein disorder was established with sequence composition being viewed as a function of IDR length (Radivojac, 2004), determination method (Garner et al., 1998), or being highly biased in certain functional sites, such as histone tails (Hansen et al., 2006) of polyQ regions (Totzeck et al., 2017). In light of the presented results, the capacity for binding and the structural nature of the binding partner have equally deep influence on sequence composition (Figure 1). Most notably, while IDPs in general are in fact depleted in hydrophobic residues, IDPs forming complexes via mutual synergistic folding are prime exceptions to this rule. Protein stability requires a hydrophobic core that stabilizes tertiary structure, and for IDPs binding to globular domains, the core is already provided by the partner, hence the interacting IDP does not have to contribute to the overall tertiary stability. However, when all interactors are disordered, the hydrophobic content must reside in these sequences, as hydrophobic collapse happens during the binding event.

The dependence of IDP features on the binding partner is also represented in their bound structures. Although helical binding (Cheng et al., 2007) and beta-augmentation (Remaut and Waksman, 2006) represent possibly the two most well-known binding modes in coupled folding and binding, analysis of the complete available set of such complexes show that such IDP segments overwhelmingly prefer irregular conformations in their bound forms. This possibly mirrors the uneven roles of the proteins in the interaction; as the folding of the ordered partner is already complete by the time of the interaction, the IDP partner has to adapt to the presented sterical constraints. In contrast, IDPs involved in synergistic folding form the core together and exhibit certain characteristics of ordered structures, most notably the relative stability of the interaction, which relies significantly on intrachain interactions. These uncovered structural differences mirror the sequential characteristics of the respective interaction classes. However, in contrast to sequence, the bound structure is strikingly more characteristic of these classes; while certain sequence compositions are compatible with multiple modes of binding, the three types of interactions are very clearly separated in the structural space.

The overlap between the three types of interactions is the most pronounced at the functional level. Most high-level biological functions arise through a deeply interconnected network of interactions of all types (Figure 3). Considering more specific processes however, reveals a generic trend showing that the importance of IDP mediated interactions increase for processes closer to the DNA. This is shown at the functional level, with DNA-, transcription-, and gene expression-related functions being enriched in IDP interactions. In addition, this trend also becomes apparent when focusing on subcellular localization (Figure 4), with ordered interactions dominating the extracellular space and the cytosol, while IDP interactions enriched in the nucleus. Furthermore, this ‘disorder-attraction’ of the DNA also differentiates between IDP-ordered and IDP-IDP interactions, with the former pertaining to DNA regulation and the latter to DNA information content. This discrimination is also reflected in the increased importance of methylation in the regulation of synergistically folding complexes (Figure 6), with methylation being primarily connected to the information access control of DNA.

As IDP-mediated functions are critically important in the cell, especially for the maintenance and processing of genetic material, the interactions involved are very precisely controlled. These control mechanisms have been studied in detail mainly for interactions arising via coupled folding and binding, especially in the context of linear motifs (Van Roey et al., 2014; Weatheritt et al., 2012b). Our study shows that various regulatory mechanisms – most notably post-translational modifications (PTMs) – also heavily modulate interactions mediated solely by IDPs in a highly coordinated but structurally indirect fashion (Figure 6). Analysis of specific examples shed light on how various PTMs affect protein function. Figure 8A shows the basic effects of PTMs for synergistically folding complexes. PTMs on one hand can directly modulate the interaction by operating as a binary on/off switch (Proctor et al., 2011), assisting partner selection (Kang et al., 2011), or fine-tuning the affinity (Teufel et al., 2009). However, PTMs can affect function indirectly by not modifying the mutually folded complex, as in the case of the Max dimeric transcriptional repressor. In this case the phosphorylation state of Max determines its DNA-binding capacity (Koskinen et al., 1994). An even more indirect modulation of function is displayed for the retinoblastoma protein Rb. In resting phase, Rb binds to the E2F1/DP1 transcription factor complex, inhibiting the transcription of genes needed for the G1/S transition. This repressor function is enhanced by the methylation of Rb on K860, recruiting L3MBTL1 (Saddic et al., 2010). L3MBTL1 is a direct repressor of transcription via chromatin compaction, augmenting the effect of Rb through a related but separate mechanism extrinsic to the Rb/E2F1/DP1 complex.

**Figure 8:**
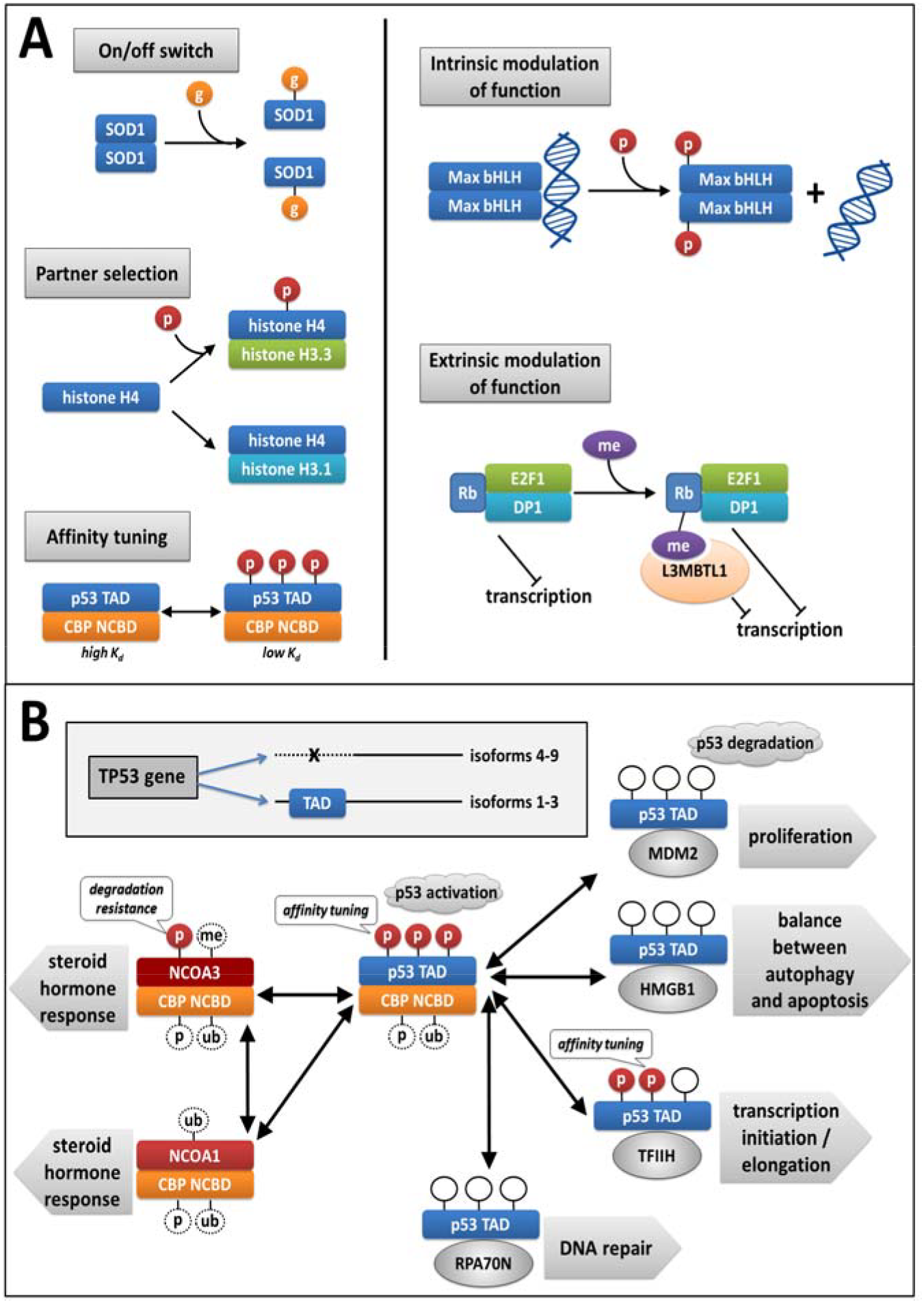
Regulatory mechanisms of complexes with mutual synergistic folding. A: examples of regulation and function modulation through post-translational modifications. p - phosphorylation, g - glutathionylation, me - methylation, SOD - superoxide dismutase, CBP - CREB-binding protein, Rb - retinoblastoma associated protein. B: interactions and their regulation in the p53/CBP regulatory sub-network. Color bars represent IDPs, grey ovals represent ordered proteins. Dashed circles represent PTMs with unknown functional effects identified in high-throughput measurements.

Apart from the modulation of individual interactions, PTMs along with other regulatory mechanisms assist the assembly of interaction networks formed by interactions from various classes. This is best exemplified by the p53/CBP regulatory sub-network that lies at the intersection of a range of critical regulatory and signaling processes. p53 is the main tumor suppressor in multicellular organisms capable of initiating apoptosis upon irreparable DNA-damage (Bieging et al., 2014). Human p53 has 9 isoforms, formed by alternative splicing, three of which (including the canonical sequence) contains an N-terminal transactivation domain (TAD). The activation of p53 depends critically on the interaction of TAD with CBP. However, this interaction is mutually exclusive with a range of other interactions between p53 TAD and ordered proteins, the most important of which is MDM2 (Ferreon et al., 2009). MDM2 is a ubiquitin ligase, the main conduit for p53 degradation. This way, competition between MDM2 and CBP for p53 is the main contributor to the control of turnover and activity of p53. The competition between MDM2, CBP and several other p53-binding proteins however, is heavily modulated by three PTM sites on p53 TAD that tune the binding affinities (Jenkins et al., 2012; Okuda and Nishimura, 2014). In addition, the p53/CBP interaction is also modulated by the same CBP region being able to bind to two members of the nuclear receptor coactivator (NCOA) family forming mutually exclusive interactions. In turn, both NCOA regions are also controlled via PTMs, e.g. conferring degradation resistance (Yi et al., 2008) that further modulates the balance in competition between binding events. The complexity of this intricate network of interactions reflects the biological complexity presented by a wide range of intertwined processes, including the regulation of proliferation, apoptosis, autophagy, DNA repair and various hormone responses.

While the majority of mutually folding IDPs can be efficiently grouped based on their sequential and structural features, a small but biologically equally important subset of them, including interactions of the p53/CBP sub-network, defy this classification scheme (Figure 7). The only unifying features of these interactions is the heavy involvement of regulatory mechanisms and their prevalence in non-membrane bounded organelles. Emerging realization of the importance of IDPs in liquid phase separation underlines the future targeted research of this highly heterogeneous group, along with the study of the five canonical classes described (Boeynaems et al., 2017; Brangwynne et al., 2015).

The uncovered differences between various types of interactions in terms of sequence, structure, function and regulation present the first step in basic understanding of how the interplay between protein folding and interaction modulates critical properties of the resulting complexes. This understanding will hopefully contribute to the ignition of the targeted research of the previously unexplored regions of the protein interactome, leading to the development of novel biomedical targeting efforts. While this might seem far-fetched at the time, not so long ago IDPs were generally considered pharmaceutically untargetable. The basic structural understanding of coupled folding and binding events, coupled with an ever increasing interest in cancer therapeutics, however, soon led to the successful clinical use of small molecule inhibitors of the p53-MDM2 interaction (Shen and Maki, 2011). While there are no current available treatments targeting mutual synergistic folding, let’s all hope (in this case) that history repeats itself.

## Acknowledgements

B.M. is the recipient of the postdoctoral fellowship of the Hungarian Academy of Sciences. L.D. is supported by the UNKP-17-3 new national excellence program of the ministry of human capacities. G.E.T. and L.D. are recipients of grant K119287 from the Hungarian Scientific Research Fund (OTKA) and “Momentum” Program of the Hungarian Academy of Sciences (LP2012/35). L.D., G.E.T. and I.S. are supported by project no. FIEK_16-1-2016-0005 financed under the FIEK_16 funding scheme (National Research, Development and Innovation Fund of Hungary). I.S. is the recipient of the Hungarian Research and Developments Fund OTKA K115698. Z.D. receives funding from the “Momentum” grant from the Hungarian Academy of Sciences (LP2014-18) and the OTKA grant (K108798).

## Methods

### Sequence and interaction datasets

Complexes formed by coupled folding and binding, and mutual synergistic folding were downloaded from the DIBS (Schad et al., 2017) and the MFIB (Fichó et al., 2017) databases, containing 773 and 205 protein complexes, respectively.

Complexes formed by ordered proteins were taken from the PDB by selecting structures containing dimeric protein interactions, as evidenced by the number of proteins (considering biomatrix transformations), PISA records, and the authors’ manual assignations. Two protein chains were considered to be in interaction if they have at least 5 atom pairs in contact. Only those structures were kept that did not contain any non-protein entities and where both interacting proteins consist of a single domain without any fragments, as defined by CATH (Pearl et al., 2003). The complete list of the 691 interacting ordered proteins is included in Table S1 and Table S2.

Sequences of IDPs devoid of interacting regions were generated from DisProt (Piovesan et al., 2017) records by removing sequence regions that are present in either DIBS or MFIB. Remaining sequences shorter than 5 residues were removed. The resulting set of 1,045 sequence regions is shown in Table S1. The human proteome containing 71,567 protein sequences was downloaded from UniProt (Apweiler et al., 2004) on 11 Aug. 2017.

### Sequence features

After considering various type of classifications, we found that the following amino acid categories are the most descriptive for distinguishing protein groups: hydrophobic (A, I, L, M, V), aromatic (F, W, Y), polar (N, Q, S, T), charged (H, K, R, D, E), rigid (P), flexible (G), and covalently interacting (C). Average content and standard variances for all 20 amino acids measured in various protein groups supporting this classification is shown in Figure S1. All protein sequence compositions were calculated on the reduced alphabet. When comparing proteins from the three interaction classes, compositions were calculated for one protein alone. In the classification of complexes of mutual synergistic folding, compositions were calculated for the entire complex. In these cases an 8th sequence parameter was used defined as: 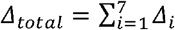 where Δ_*i*_ is the largest composition difference of residue group i between constituent chains.

### Structure features

Structural features of proteins were calculated from their bound structures. Secondary structure assignment was performed by DSSP (Touw et al., 2015) using a three-state classification distinguishing helical ('H','G','I'), extended ('B','E') and irregular ('S','T', unassigned) residues.

Molecular surfaces were calculated using Naccess (Hubbard and Thornton, 1992). Solvent accessible surface area (SASA) was defined by the Nacces absolute surface column. Interface is defined as the increase in SASA as a result of removing interaction partners from the structure. Buried surface was calculated by subtracting interface area and SASA from the sum of standard surfaces of residues in the protein chain. Thus, interface and buried surfaces represent the area that is made inaccessible to the solvent by the partner(s) or by the analysed protein itself. All calculated areas were split into hydrophobic (H) and polar (P) contributions based on the polarity of the corresponding atom. Polar/hydrophobic assignations were taken from Naccess.

Contacts were defined at the atomic level. Two atoms were considered to be in contact if their distances are shorter than the sum of the two atoms’ van der Waals radii plus 1 Angstrom.

Interaction energies for residues were calculated using the statistical potentials described in (Dosztányi et al., 2005). These interaction potentials were demonstrated to adequately describe the energetic features of interacting proteins, including IDPs (Mészáros et al., 2007).

When comparing proteins from the three interaction classes, structural parameters were calculated for one protein alone. In the classification of complexes of mutual synergistic folding, these were calculated for the entire complex.

### Functional annotations

Biological functions and subcellular localizations were taken from the DIBS and MFIB databases in the forms of GO terms. Annotations for ordered complexes were generated from the GO annotations of constituent proteins (taken from UniProt-GOA) using the approach described in DIBS/MFIB (<u>http://dibs.enzim.ttk.mta.hu/help.php</u>).

PPI GO Slim and CellLoc GO Slim were created manually from the ‘biological process’ and ‘cellular localization’ namespaces of GO, by selecting terms that are either assigned to studied complexes or are ancestors of such terms. PPI GO Slim was partitioned into two levels, with the 'Generic' part containing high level cellular/organismal processes such as 'transport', 'communication' or 'development'. In addition, the 'Specific' part of PPI GO Slim contains terms describing specific biological subprocesses, such as 'gene expression regulation', 'organelle organization' or 'proteolysis', through which generic processes are executed. The terms contained in PPI GO Slim and CellLoc GO Slim are shown in Table S3.

### Heterogeneity

Principal Component Analysis (PCA) and hierarchical and K-means clustering was done using the R statistical computing environment (version 3.3.1) (Tierney, 2012). For PCA calculations, proteins from the three interaction classes were represented by sequence and structure features defined in Table S1 and Table S2, respectively. Functional annotations were represented by a 23-element vector, where each element marks the number of GO terms that can be mapped to each of the 23 generic cellular/organismal processes of the GO PPI Slim (see Table S3). Biplots for the 7 and 11 sequences/structure parameters are shown in Figure S3 and Figure S4.

For clustering, the same sequence/structure features were used as input for the Ward.2 algorithm in R, using Euclidean distances. Dissimilarity of two proteins *i* and *j* is defined as: 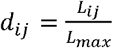, where *L*_*ij*_ is the linkage distance given by the clustering (of all proteins from all three classes) between the two proteins from the same interaction class, and *L*_*max*_ is the maximal linkage distance. Heterogeneity values are defined as the geometrical averages of dissimilarity values between all protein pairs from a given class.

In the case of functional heterogeneity, the hierarchical cluster tree was replaced by the GO ontology tree. Distances between terms that are in a parent/child relationship was defined as 1. Dissimilarity between two complexes was defined based on their most similar GO term pairs. Let *t*_*i*_ be the GO biological process terms of complex A and *t*_*j*_ be the GO biological process terms of complex B. For each *t*_*i*_ we choose a *t*_*j*_ pair, for which their distances in the ontology is minimal. I.e. let t* be the most specific (low level) term in the ontology that is the common parent of both *t*_*i*_ and *t*_*j*_. The distance between *t*_*i*_ and *t*_*j*_ is the distance between *t*_*i*_ and *t\**, plus the distance between *t*_*j*_ and *t\**. Next, we normalize this distance with the maximal possible distance that could be between *t*_*i*_ and *t*_*j*_, i.e. the sum of the distances of the two terms and the ontology root (‘biological_process’). The dissimilarity between two complexes in the functional sense is defined as the average normalized distance between their term pairs, selected for minimal distance. From these measures, heterogeneity values are derived in the same fashion as for sequence and structure, described above.

### Regulation

PTMs were taken from the 2 October 2017 version of PhosphoSitePlus (Hornbeck et al., 2015), and were mapped to complex structures using BLAST between UniProt and PDB sequences. Protein isoforms were taken from UniProt. To determine alternative binding partners for IDPs, all oligomer PDB structure containing the same UniProt region were selected. PDB structures listed as related in the corresponding MFIB or DIBS entry were removed. Structures containing the same interaction partners as the original complex were also removed. For data compiled regarding regulatory mechanisms in the three interaction classes, see Table S4.

## References

Andreeva, A., Howorth, D., Chandonia, J.-M., Brenner, S.E., Hubbard, T.J.P., Chothia, C., and Murzin, A.G. (2008). Data growth and its impact on the SCOP database: new developments. Nucleic Acids Res. 36, D419–D425.

Apweiler, R., Bairoch, A., Wu, C.H., Barker, W.C., Boeckmann, B., Ferro, S., Gasteiger, E., Huang, H., Lopez, R., Magrane, M., et al. (2004). UniProt: the Universal Protein knowledgebase. Nucleic Acids Res. 32, D115–D119.

Bah, A., and Forman-Kay, J.D. (2016). Modulation of Intrinsically Disordered Protein Function by Post-translational Modifications. J. Biol. Chem. 291, 6696–6705.

Bieging, K.T., Mello, S.S., and Attardi, L.D. (2014). Unravelling mechanisms of p53-mediated tumour suppression. Nat. Rev. Cancer 14, 359–370.

Boeynaems, S., Bogaert, E., Kovacs, D., Konijnenberg, A., Timmerman, E., Volkov, A., Guharoy, M., De Decker, M., Jaspers, T., Ryan, V.H., et al. (2017). Phase Separation of C9orf72 Dipeptide Repeats Perturbs Stress Granule Dynamics. Mol. Cell 65, 1044–1055.e5.

Brangwynne, C.P., Tompa, P., and Pappu, R.V. (2015). Polymer physics of intracellular phase transitions. Nat. Phys. 11, 899–904.

Buljan, M., Chalancon, G., Dunker, A.K., Bateman, A., Balaji, S., Fuxreiter, M., and Babu, M.M. (2013). Alternative splicing of intrinsically disordered regions and rewiring of protein interactions. Curr. Opin. Struct. Biol. 23, 443–450.

Campen, A., Williams, R.M., Brown, C.J., Meng, J., Uversky, V.N., and Dunker, A.K. (2008). TOP-IDP-scale: a new amino acid scale measuring propensity for intrinsic disorder. Protein Pept. Lett. 15, 956–963.

Cheng, Y., Oldfield, C.J., Meng, J., Romero, P., Uversky, V.N., and Keith Dunker, A. (2007). Mining α-Helix-Forming Molecular Recognition Features with Cross Species Sequence Alignments†. Biochemistry 46, 13468–13477.

Chu, X., and Wang, J. (2014). Specificity and affinity quantification of flexible recognition from underlying energy landscape topography. PLoS Comput. Biol. 10, e1003782.

Demarest, S.J., Martinez-Yamout, M., Chung, J., Chen, H., Xu, W., Dyson, H.J., Evans, R.M., and Wright, P.E. (2002). Mutual synergistic folding in recruitment of CBP/p300 by p160 nuclear receptor coactivators. Nature 415, 549–553.

Deng, X., Gumm, J., Karki, S., Eickholt, J., and Cheng, J. (2015). An Overview of Practical Applications of Protein Disorder Prediction and Drive for Faster, More Accurate Predictions. Int. J. Mol. Sci. 16, 15384–15404.

Dosztányi, Z., Csizmók, V., Tompa, P., and Simon, I. (2005). The pairwise energy content estimated from amino acid composition discriminates between folded and intrinsically unstructured proteins. J. Mol. Biol. 347, 827–839.

Dosztányi, Z., Mészáros, B., and Simon, I. (2010). Bioinformatical approaches to characterize intrinsically disordered/unstructured proteins. Brief. Bioinform. 11, 225–243.

Dyson, H.J., and Wright, P.E. (2002). Coupling of folding and binding for unstructured proteins. Curr. Opin. Struct. Biol. 12, 54–60.

Dyson, H.J., and Wright, P.E. (2005). Intrinsically unstructured proteins and their functions. Nat. Rev. Mol. Cell Biol. 6, 197–208.

Ferreon, J.C., Lee, C.W., Arai, M., Martinez-Yamout, M.A., Dyson, H.J., and Wright, P.E. (2009). Cooperative regulation of p53 by modulation of ternary complex formation with CBP/p300 and HDM2. Proc. Natl. Acad. Sci. U. S. A. 106, 6591–6596.

Fichó, E., Reményi, I., Simon, I., and Mészáros, B. (2017). MFIB: a repository of protein complexes with mutual folding induced by binding. Bioinformatics.

Garner, E., Cannon, P., Romero, P., Obradovic, Z., and Dunker, A.K. (1998). Predicting Disordered Regions from Amino Acid Sequence: Common Themes Despite Differing Structural Characterization. Genome Inform. Ser. Workshop Genome Inform. 9, 201–213.

Gast, K., Damaschun, H., Eckert, K., Schulze-Forster, K., Maurer, H.R., Müller-Frohne, M., Zirwer, D., Czarnecki, J., and Damaschun, G. (1995). Prothymosin alpha: a biologically active protein with random coil conformation. Biochemistry 34, 13211–13218.

Gsponer, J., Futschik, M.E., Teichmann, S.A., and Babu, M.M. (2008). Tight regulation of unstructured proteins: from transcript synthesis to protein degradation. Science 322, 1365–1368.

Gunasekaran, K., Tsai, C.-J., and Nussinov, R. (2004). Analysis of ordered and disordered protein complexes reveals structural features discriminating between stable and unstable monomers. J. Mol. Biol. 341, 1327–1341.

Hansen, J.C., Lu, X., Ross, E.D., and Woody, R.W. (2006). Intrinsic protein disorder, amino acid composition, and histone terminal domains. J. Biol. Chem. 281, 1853–1856.

Hornbeck, P.V., Zhang, B., Murray, B., Kornhauser, J.M., Latham, V., and Skrzypek, E. (2015). PhosphoSitePlus, 2014: mutations, PTMs and recalibrations. Nucleic Acids Res. 43, D512–D520.

Hsu, W.-L., Oldfield, C.J., Xue, B., Meng, J., Huang, F., Romero, P., Uversky, V.N., and Dunker, A.K. (2013). Exploring the binding diversity of intrinsically disordered proteins involved in one-to-many binding. Protein Sci. 22, 258–273.

Hubbard, S., and Thornton, J. (1992). Naccess.

Jenkins, L.M.M., Durell, S.R., Mazur, S.J., and Appella, E. (2012). p53 N-terminal phosphorylation: a defining layer of complex regulation. Carcinogenesis 33, 1441–1449.

Kang, B., Pu, M., Hu, G., Wen, W., Dong, Z., Zhao, K., Stillman, B., and Zhang, Z. (2011). Phosphorylation of H4 Ser 47 promotes HIRA-mediated nucleosome assembly. Genes Dev. 25, 1359–1364.

Koskinen, P.J., Västrik, I., Mäkelä, T.P., Eisenman, R.N., and Alitalo, K. (1994). Max activity is affected by phosphorylation at two NH2-terminal sites. Cell Growth Differ. 5, 313–320.

Mackinnon, S.S., Malevanets, A., and Wodak, S.J. (2013). Intertwined associations in structures of homooligomeric proteins. Structure 21, 638–649.

Malhis, N., Jacobson, M., and Gsponer, J. (2016). MoRFchibi SYSTEM: software tools for the identification of MoRFs in protein sequences. Nucleic Acids Res. 44, W488–W493.

Meng, F., Uversky, V.N., and Kurgan, L. (2017). Comprehensive review of methods for prediction of intrinsic disorder and its molecular functions. Cell. Mol. Life Sci. 74, 3069–3090.

Mészáros, B., Tompa, P., Simon, I., and Dosztányi, Z. (2007). Molecular principles of the interactions of disordered proteins. J. Mol. Biol. 372, 549–561.

Mészáros, B., Simon, I., and Dosztányi, Z. (2009). Prediction of protein binding regions in disordered proteins. PLoS Comput. Biol. 5, e1000376.

Monastyrskyy, B., Kryshtafovych, A., Moult, J., Tramontano, A., and Fidelis, K. (2014). Assessment of protein disorder region predictions in CASP10. Proteins 82 Suppl 2, 127–137.

Okuda, M., and Nishimura, Y. (2014). Extended string binding mode of the phosphorylated transactivation domain of tumor suppressor p53. J. Am. Chem. Soc. 136, 14143–14152.

Pearl, F.M.G., Bennett, C.F., Bray, J.E., Harrison, A.P., Martin, N., Shepherd, A., Sillitoe, I., Thornton, J., and Orengo, C.A. (2003). The CATH database: an extended protein family resource for structural and functional genomics. Nucleic Acids Res. 31, 452–455.

Piovesan, D., Tabaro, F., Mičetić, I., Necci, M., Quaglia, F., Oldfield, C.J., Aspromonte, M.C., Davey, N.E., Davidović, R., Dosztányi, Z., et al. (2017). DisProt 7.0: a major update of the database of disordered proteins. Nucleic Acids Res. 45, D1123–D1124.

Proctor, E.A., Ding, F., and Dokholyan, N.V. (2011). Structural and thermodynamic effects of post-translational modifications in mutant and wild type Cu, Zn superoxide dismutase. J. Mol. Biol. 408, 555–567.

Radivojac, P. (2004). Protein flexibility and intrinsic disorder. Protein Sci. 13, 71–80.

Redfern, O.C., Dessailly, B., and Orengo, C.A. (2008). Exploring the structure and function paradigm. Curr. Opin. Struct. Biol. 18, 394–402.

Remaut, H., and Waksman, G. (2006). Protein-protein interaction through beta-strand addition. Trends Biochem. Sci. 31, 436–444.

Richmond, T.J. (1984). Solvent accessible surface area and excluded volume in proteins. Analytical equations for overlapping spheres and implications for the hydrophobic effect. J. Mol. Biol. 178, 63–89.

Rumfeldt, J.A.O., Galvagnion, C., Vassall, K.A., and Meiering, E.M. (2008). Conformational stability and folding mechanisms of dimeric proteins. Prog. Biophys. Mol. Biol. 98, 61–84.

Saddic, L.A., West, L.E., Aslanian, A., Yates, J.R., 3rd, Rubin, S.M., Gozani, O., and Sage, J. (2010). Methylation of the retinoblastoma tumor suppressor by SMYD2. J. Biol. Chem. 285, 37733–37740.

Schad, E., Fichó, E., Pancsa, R., Simon, I., Dosztányi, Z., and Mészáros, B. (2017). DIBS: a repository of disordered binding sites mediating interactions with ordered proteins. Bioinformatics.

Shen, H., and Maki, C.G. (2011). Pharmacologic activation of p53 by small-molecule MDM2 antagonists. Curr. Pharm. Des. 17, 560–568.

Sutovsky, H., and Gazit, E. (2004). The von Hippel-Lindau tumor suppressor protein is a molten globule under native conditions: implications for its physiological activities. J. Biol. Chem. 279, 17190–17196.

Teufel, D.P., Bycroft, M., and Fersht, A.R. (2009). Regulation by phosphorylation of the relative affinities of the N-terminal transactivation domains of p53 for p300 domains and Mdm2. Oncogene 28, 2112–2118.

Tierney, L. (2012). The R Statistical Computing Environment. In Lecture Notes in Statistics, pp. 435–447.

Tompa, P. (2011). Unstructural biology coming of age. Curr. Opin. Struct. Biol. 21, 419–425.

Tompa, P., and Fuxreiter, M. (2008). Fuzzy complexes: polymorphism and structural disorder in protein-protein interactions. Trends Biochem. Sci. 33, 2–8.

Totzeck, F., Andrade-Navarro, M.A., and Mier, P. (2017). The Protein Structure Context of PolyQ Regions. PLoS One 12, e0170801.

Touw, W.G., Baakman, C., Black, J., te Beek, T.A.H., Krieger, E., Joosten, R.P., and Vriend, G. (2015). A series of PDB-related databanks for everyday needs. Nucleic Acids Res. 43, D364–D368.

Tsai, C.J., Kumar, S., Ma, B., and Nussinov, R. (1999). Folding funnels, binding funnels, and protein function. Protein Sci. 8, 1181–1190.

Uversky, V.N., Gillespie, J.R., and Fink, A.L. (2000). Why are “natively unfolded” proteins unstructured under physiologic conditions? Proteins 41, 415–427.

Van Roey, K., Uyar, B., Weatheritt, R.J., Dinkel, H., Seiler, M., Budd, A., Gibson, T.J., and Davey, N.E. (2014). Short linear motifs: ubiquitous and functionally diverse protein interaction modules directing cell regulation. Chem. Rev. 114, 6733–6778.

Wang, N., Majmudar, C.Y., Pomerantz, W.C., Gagnon, J.K., Sadowsky, J.D., Meagher, J.L., Johnson, T.K., Stuckey, J.A., Brooks, C.L., 3rd, Wells, J.A., et al. (2013). Ordering a dynamic protein via a small-molecule stabilizer. J. Am. Chem. Soc. 135, 3363–3366.

Weatheritt, R.J., Luck, K., Petsalaki, E., Davey, N.E., and Gibson, T.J. (2012a). The identification of short linear motif-mediated interfaces within the human interactome. Bioinformatics 28, 976–982.

Weatheritt, R.J., Davey, N.E., and Gibson, T.J. (2012b). Linear motifs confer functional diversity onto splice variants. Nucleic Acids Res. 40, 7123–7131.

Wright, P.E., and Dyson, H.J. (1999). Intrinsically unstructured proteins: re-assessing the protein structure-function paradigm. J. Mol. Biol. 293, 321–331.

Wright, P.E., and Dyson, H.J. (2015). Intrinsically disordered proteins in cellular signalling and regulation. Nat. Rev. Mol. Cell Biol. 16, 18–29.

Yi, P., Feng, Q., Amazit, L., Lonard, D.M., Tsai, S.Y., Tsai, M.-J., and O’Malley, B.W. (2008). Atypical protein kinase C regulates dual pathways for degradation of the oncogenic coactivator SRC-3/AIB1. Mol. Cell 29, 465–476.

